# Volumetric Lissajous Confocal Microscopy

**DOI:** 10.1101/735654

**Authors:** Takahiro Deguchi, Paolo Bianchini, Gemma Palazzolo, Michele Oneto, Alberto Diaspro, Martí Duocastella

**Affiliations:** Nanoscopy & NIC@IIT, Istituto Italiano di Tecnologia, via Morego 30, 16163 Genova, Italy; Enhanced Regenerative Medicine, Istituto Italiano di Tecnologia, via Morego 30, 16163 Genova, Italy; Dipartimento di Fisica, Universita di Genova, Via Dodecaneso 33, 16146, Genoa, Italy

## Abstract

Dynamic biological systems present challenges to existing three-dimensional (3D) optical microscopes because of their continuous temporal and spatial changes. Most techniques are based on rigid architectures, as in confocal microscopy, where a laser beam is sequentially scanned at a predefined spatial sampling rate and pixel dwell time. Here, we developed volumetric Lissajous confocal microscopy to achieve unsurpassed 3D scanning speed with a tunable sampling rate. The system combines an acoustic liquid lens for continuous axial focus translation with a resonant scanning mirror. Accordingly, the excitation beam follows a dynamic Lissajous trajectory enabling sub-millisecond acquisitions of image series containing 3D information at a sub-Nyquist sampling rate. By temporal accumulation and/or advanced interpolation algorithms, volumetric imaging rate is selectable using a post-processing step at the desired spatiotemporal resolution for events of interest. We demonstrate multicolor and calcium imaging over volumes of tens of cubic microns with acquisition speeds up to 5 kHz.

## 1. Introduction

Confocal laser scanning microscopy (CLSM) has arguably become the *de facto* standard for the three-dimensional (3D) characterization of biological processes at the sub-cellular level [1–4]. Based on the raster scanning of a laser beam for illumination CLSM provides optical sectioning with synchronous multi-channel imaging. In addition, these confocal architectures are compatible with super-resolution microscopy and advanced spectroscopic techniques such as fluorescence correlation spectroscopy and fluorescence lifetime imaging [5–9]. However, despite the complete toolset that CLSM represents, a central question in this imaging modality is how to select the main scanning parameters (number of pixels, pixel dwell time and scanned volume) to maximize the spatiotemporal information retrieved from a sample. While the answer is straightforward when characterizing fixed samples or slowly varying processes — we can always use long pixel dwell times and a large number of pixels to attain high signal to noise ratio (SNR) and spatial resolution — problems arise when imaging fast phenomena or rapidly evolving systems. In these time-sensitive cases, one is typically faced with the dilemma of having to sacrifice either spatial or temporal resolution. This is common in most imaging technologies — reducing the number of sampled points increases imaging speed, albeit with a loss of spatial resolution — but in CLSM there are two aspects that further aggravate such a tradeoff. First, current confocal systems are unsuitable for fast 3D scanning. By using resonant mirrors lateral (X,Y) beam scanning at microsecond timescales can be routinely achieved [10], but axial focus translation (Z) remains one or two orders of magnitude slower [11]. Thus 3D sampling is unevenly performed — fast along XY, slow along Z — greatly limiting volumetric imaging rates. Secondly, even if a desired sampling rate and optimal tradeoff between imaging speed and resolution can be selected, it must remain fixed for the entirety of an acquisition. In other words, the rigid scanning architecture of confocal microscopes impedes adapting the scanning parameters as samples evolve over time. As a result, confocal microscopes remain sub-optimal tools to investigate the dynamics of fast processes or multiscale systems that occur over different temporal scales.

Volumetric Lissajous confocal microscopy addresses the limitations encountered with existing CLSM systems. By integrating an ultrasound-driven liquid lens (TAG lens) into a confocal workstation equipped with a resonant mirror, we achieve three-dimensional beam scanning at kHz rates. This high-speed comes with the caveat of sub-optimal sampling — the scan trajectories correspond to dynamic Lissajous patterns that sample only a fraction of the voxels within a volume. However, by properly selecting the TAG lens driving frequency the trajectories can be rendered dynamic, enabling trajectory variations over time. As a result, each volumetric scan samples different voxels and by accumulating multiple scans, the spatial resolution of the reconstructed 3D image can be improved. This represents a paradigm shift in laser scanning microscopy techniques — information is continuously acquired, and a post-processing step allows the volumetric imaging rate to be selected based on the desired spatial or temporal resolution. In addition, Lissajous scanning is compatible with inpainting algorithms used to fill the missing information in undersampled images, helping to restore spatial resolution without sacrificing time. We present a detailed description and characterization of the technique, and demonstrate its potential for fast 3D live cell imaging by capturing the calcium signals of neuronal networks from an *in vitro* brain model.

## 2. Principle and implementation of Volumetric Lissajous confocal microscopy

Our approach uses two harmonically oscillating systems to achieve high-speed 3D scanning. A liquid lens driven by ultrasound (TAG lens) at ~456 kHz is used for z-focus scanning, and a resonant mirror at ~8 kHz for x-scanning. Volumetric scanning is completed with a galvo mirror system to linearly translate the beam along the y-axis (slow axis in our microscope). As shown in Figure 1a, the resulting 3D scan trajectories correspond to Lissajous patterns exhibiting two key features. First, a large increase in laser scanning speed is achieved, up to 45-fold faster for our experimental conditions, compared with conventional scanning methods with a single resonant mirror (see Supplementary information and Figure S1). Even if Lissajous trajectories have been reported in different microscopy techniques, including atomic force microscopy or two-photon microscopy [12–14], they have been limited to 2D scans along the xy plane. Instead, our system extends their use into 3D imaging utilizing the superior scanning speed in the z-axis. Secondly, the Lissajous patterns can be adjusted by tuning the TAG lens driving frequency (Figure 1b). Notably, not only can the shape of the trajectory be controlled, but also its temporal characteristics enabling the generation of static or dynamic patterns. The dynamic patterns, as illustrated in Figure 1b, can increase the number of sampled voxels over multiple scans, providing a unique method to correlate spatial sampling with imaging rate.

**Figure 1.**
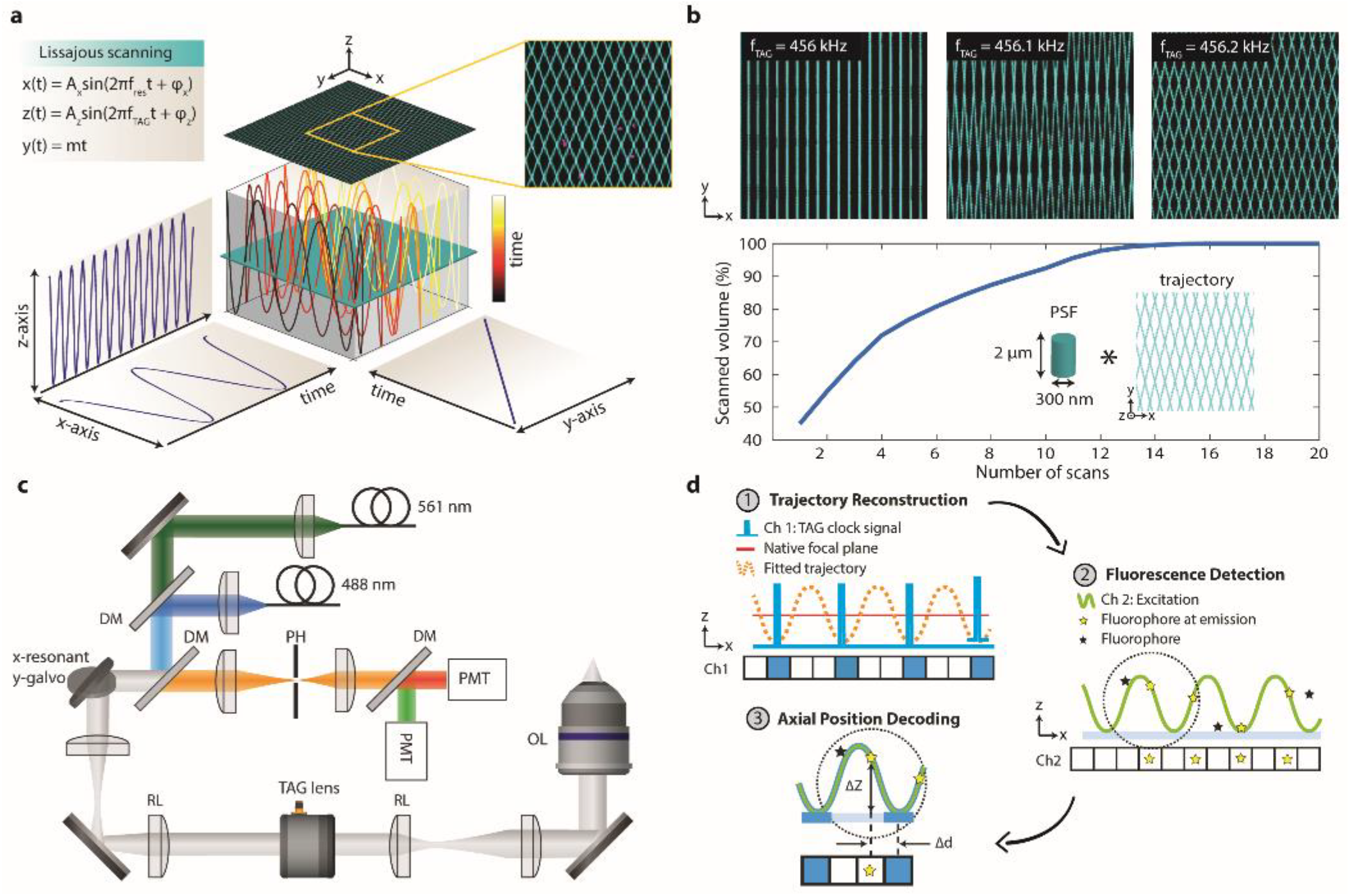
Principle and implementation of the Lissajous confocal microscope. a) Scheme of a 3D-Lissajous trajectory generated by scanning a beam using a resonant mirror (x-axis), the TAG lens (z-axis), and a galvo mirror (y-axis). The inset corresponds to an experimentally recorded trajectory obtained by placing a thin fluorescence sample in the microscope. b) Simulated trajectories considering a resonant mirror operated at 8 kHz and at various TAG lens frequencies. Plot of the fraction of scanned volume versus number of scans for fTAG = 456.2 kHz. The values were calculated by convolving a cylindrical point-spread function with simulated 3D trajectories. Because the trajectories are temporally dynamic, scanning multiple times enables increasing the number of sampled voxels. c) Scheme of the experimental setup. DM: dichroic mirror, PMT: photomultiplier tube; PH: pinhole; OL: objective lens; RL: relay lens. Only two lasers and two detectors out of four are illustrated. d) Workflow of the three-step process used to retrieve 3D images without the need of fast electronics.

Volumetric Lissajous confocal microscopy is simple to implement in any commercial confocal system with a resonant mirror. The main modification involves inserting the TAG lens at the conjugate plane of the back focal plane of the microscope objective lens. In our experiments, we placed the TAG lens and a relay lens system between the tube lens and the scan lens of the microscope, as illustrated in Figure 1c. One key aspect when implementing the Lissajous microscope is synchronizing the scanned excitation with the detection. This task is typically accomplished with fast acquisition cards [15,16]. However, the relative high frequency of the TAG lens requires electronics significantly faster than those commonly employed [16]. To ease implementation, we designed a three-step approach that obviates the need of fast electronics, as shown in Figure 1d. The Lissajous pattern is first reconstructed, line by line, by taking into account the clock signals of the TAG lens (directly fed into Channel 1 of the acquisition card of the microscope) enabling the axial position corresponding to each pixel to be defined. The fluorescence signal of the sample, simultaneously detected in Channel 2, is then analyzed and the axial information intrinsically contained for each pixel is decoded, leading to the reconstruction of a 3D image.

## 3. Results

### 3.1 Optical performance of the microscope

Initially, we characterized the optical performance of the Lissajous microscope by imaging 100 nm fluorescent nanospheres volumetrically dispersed in 1 % agarose gel using a 40x (1.15 NA) objective lens. To address any loss of spatial resolution caused by sub-optimal sampling and obtain a high SNR, we averaged the Lissajous trajectories over 250 scans. The sample information was continuously acquired and a z-stack containing 32 optical sections (this number can be arbitrarily selected) was computed using a post-processing step. Figure 2a shows the reconstructed 3D image, in which the axial position (depth) of the spheres has been color-coded. The nanospheres can be distinguished within an axial range (Δz) of 70 µm, about 30 times the native depth of field of the objective. The axial range depends on several factors such as the optical magnification of the system, the driving frequency and voltage of the TAG lens, and the objective lens used [17,18]. For example, a 20x (0.8 NA) objective lens realized a 200 µm axial range at the same imaging conditions as the 40x objective (Figure S2). By benchmarking the image against those obtained with a traditional piezo z-stage (Figure 2b) we can validate the suitability of Lissajous scans and the reconstruction process for optical microscopy. There is an excellent match between the two images with only small discrepancies at the bottom and the top of the axial range, as to be expected from aberrations induced by the TAG lens at these positions [15,18–20].

**Figure 2.**
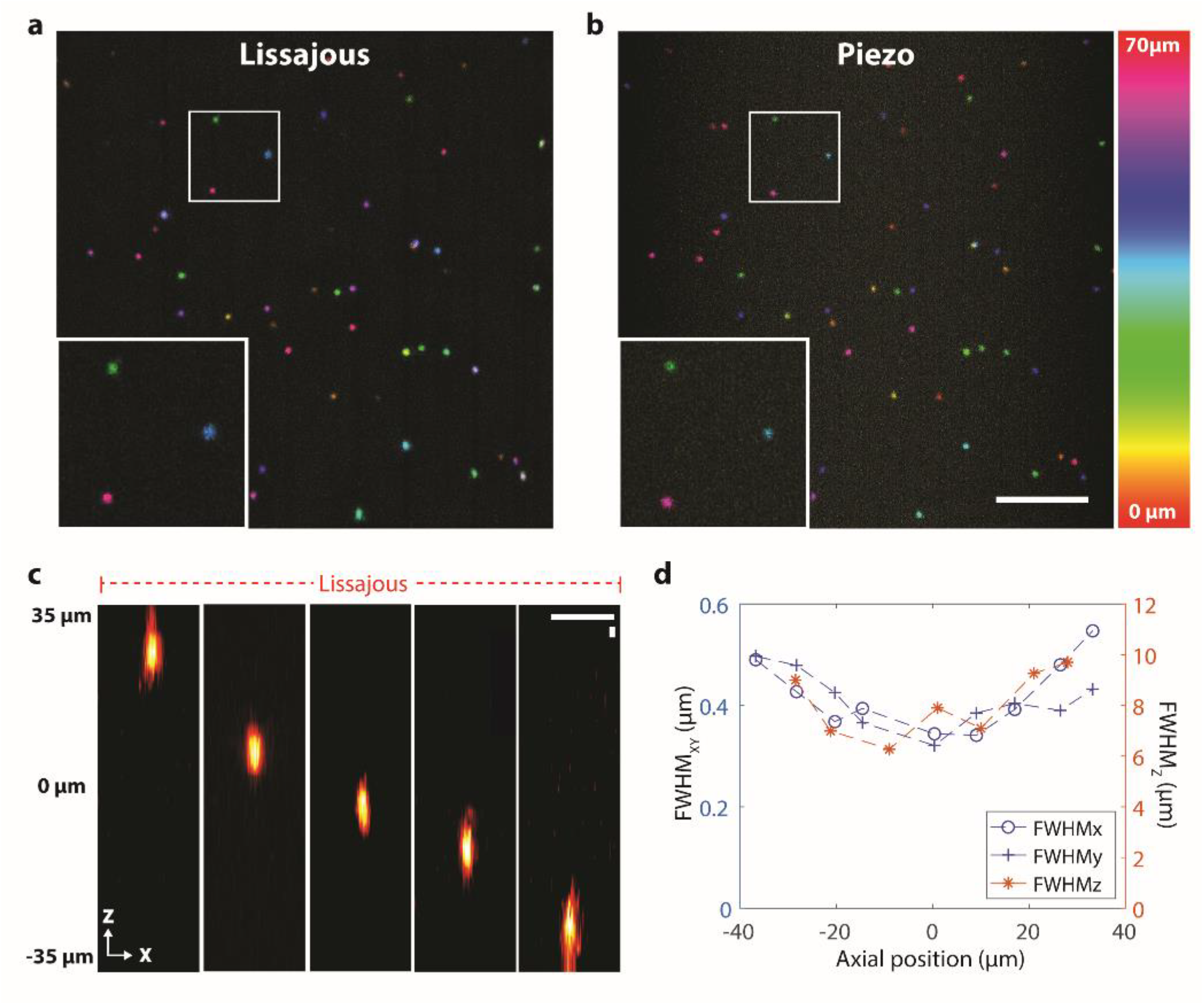
Volumetric imaging and system characterization. a) Depth-color coded images of 100 nm nanospheres in 3D agarose taken by Lissajous scan and b) standard piezo scan over a 70 µm axial range. Image acquisition time for both images were 9.3 seconds, corresponding to 280 scans. The different colors represent different depth indicated in the color bar. Scale bar 10 µm. c) Point spread function of the Lissajous microscope obtained at different axial positions for the conditions reported in a). Scale bar 2 µm. d) FWHMs of the PSFs of Lissajous scan in XYZ directions at different axial positions.

A more quantitative analysis of the spatial resolution of our microscope can be obtained by measuring its point spread function (PSF) at different focal planes using the nanospheres, as illustrated in Figure 2c. For all planes the shape of the PSF is straight and symmetrical along the optical axis, with only small distortions close to the extrema of the axial range. Indeed, the lateral and axial full width at the half maximum (FWHM) of the PSF remain practically constant within ± 20 µm of the native plane of the objective, but becomes larger beyond this range (Figure 2d). Note the clear asymmetry in spatial resolution between the x-y and z axis — while the average FWHM of the lateral PSF is about 0.4 µm, in agreement with the 0.37 µm measured with the standard piezo, the axial PSF is about 4 times longer, 8 µm vs 2.2 µm (see Supplementary information and Figure S3). The effective elongation of the PSF along the z-axis is not a fundamental limit of the technique, but rather a drawback of our simplified implementation. Since the electronic card of our commercial microscope has a limited number of time channels, for each x-scan, the collected photons are sorted into only 1024 windows or pixels. Although, in a regular confocal microscope, it is sufficient to fully reconstruct a 2D image, in our case the focus is continuously scanned along the z-axis, each time window can contain information of multiple focal planes depending on the axial range or frequency of the TAG (f_TAG_) lens (Figure S3a). Therefore, an increased number of pixel could provide the optimal pixel sampling. For instance, the conditions used here (Δz=70 µm, f_TAG_ = 457 kHz, and 40 ns time window) resulted in the merging of information of up to 4 µm along the z axis in a single pixel. For Δz = 18 µm, the integration of axial data is reduced to 1 µm, thus explaining the increase in z-resolution experimentally observed in this case (Figure S3b). Nevertheless, the optical performance of our microscope could be easily improved by decreasing the number of extrema per x-line, i.e. by employing a smaller f_TAG_ value (Figure S4) or by increasing the number of time windows using one of several commercial microscopes with resonant mirror systems capable of up to 4000 pixels per line.

Notably, our Lissajous confocal microscope preserves core advantages of confocal microscopes, including the possibility of simultaneous multi-color imaging. Figure 3 shows images of a murine brain slice captured with our system after integration over 30 scans. In this case, neurons, blood vessels and nuclei were labeled with three different dyes. The sample was simultaneously irradiated with three lasers and the corresponding information was captured with three separate point detectors. The reconstructed sections at different focal planes, over an axial range of 70 µm, and the corresponding maximum intensity projections (MIP) and 3D rendered volumes reveal an overall good image quality. The different brain components are clearly distinguished with negligible cross talk between the channels (see Supplementary Movie 1). These results also indicate that chromatic aberration effects are not significant in the current implementation, in agreement with previous works on TAG-enabled systems [15].

**Figure 3:**
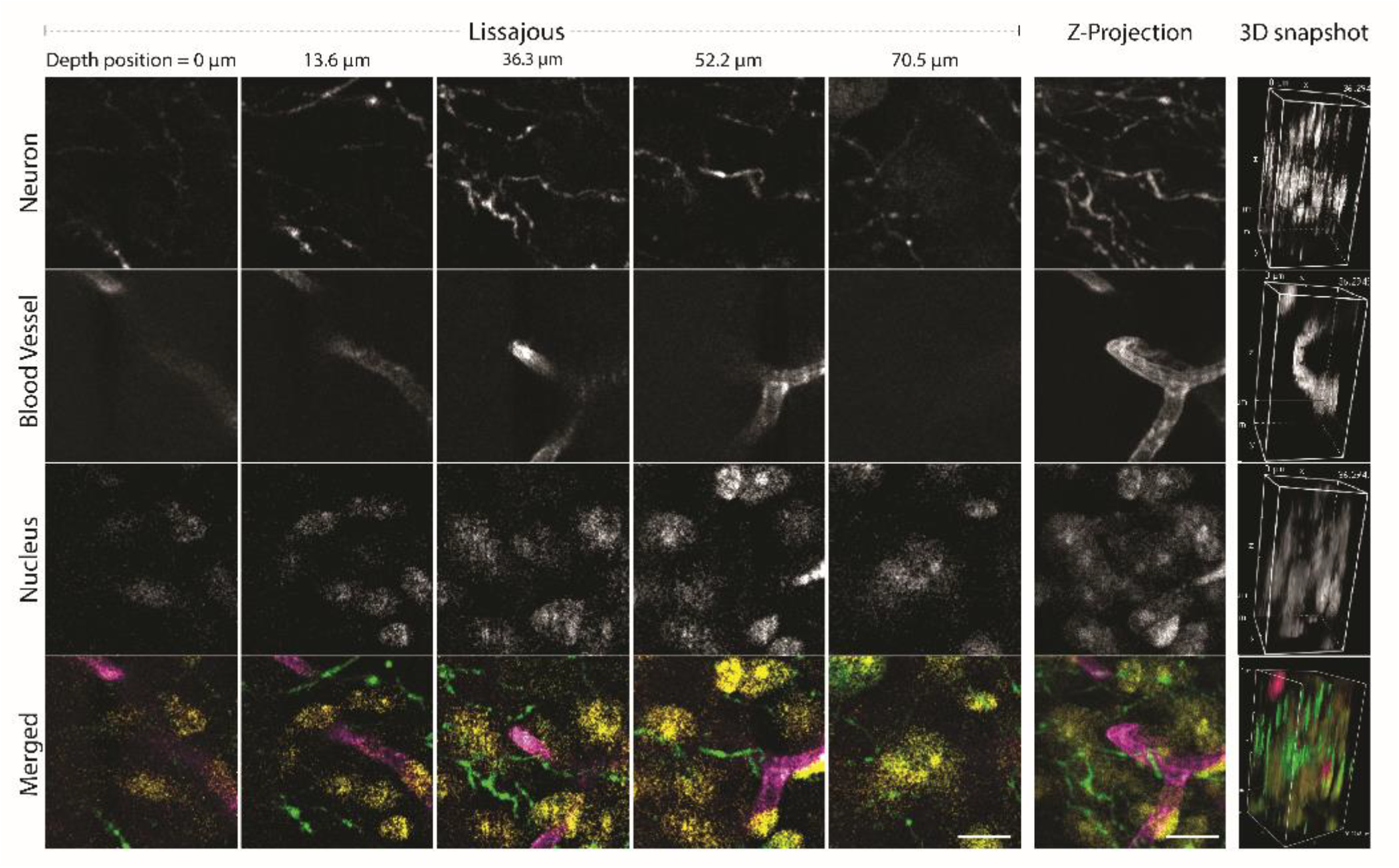
Synchronous 3-color volumetric brain imaging. Rows correspond to different color channels, each highlighting a different part of the brain. The bottom row consists of merged images containing the three channels (green for neuron, magenta for blood vessels, and yellow for nuclei). Columns indicate the z-position of the images. The last two right columns are a maximum intensity projection (MIP) and a 3D rendering. The overall axial scan range was 70.5 µm, and images were reconstructed with 30 scans. A video of the 3D rendered images is available (Movie 1). Scale bar 10 µm.

### 3.2 Instant 3D imaging with tunable spatiotemporal resolution

The key features of our microscope are 3D beam scanning at high-speed and sub-Nyquist sampling, together with time-varying spatial resolution. These two aspects are highlighted in Figure 4, where 3D images of neurons from a murine brain slice rendered by our Lissajous microscope are compared with a conventional raster scanning/piezo stage microscope. Only one 3D-scan (one frame) is needed for the Lissajous image to obtain volumetric information, with neurons clearly discernable across the entire 3D imaging space. This result is in striking contrast to the image obtained with the typical confocal system — even if the same acquisition time and laser intensity were used, only one optical section is captured. The restriction for a single acquisition is that our image does have a lower SNR and the neuronal networks appear to be disconnected due to the sparse sampling. Remarkably, however, by accumulating the number of scans the Lissajous images exhibit an increased spatial sampling and, consequently, a higher spatial resolution and SNR. Indeed, after accumulating 5 scans more neuronal processes become recognizable, and after 10 scans the neuronal networks extending in the 3D space become completely traceable. Integration over 30 scans results in images where all points have been sampled, in agreement with simulations from Figure 1b. The temporal evolution of the volumetric information retrieved with the Lissajous microscope is drastically different from that observed in a conventional confocal system. Conventional methods require sequentially capturing multiple focal planes, a time-consuming task due to the need of mechanical z-focus translation. As such acquisition is directional (from bottom to top in current experiment) and after 30 scans (or frames) only half the volume of interest has been imaged, with the remaining volume being totally unknown. This comparison between the two microscope modalities manifests the unprecedented flexibility of Lissajous scanning. As 3D structural information is immediately accessed and the accumulation of multiple scans enhances spatial resolution, scanning parameters do not need to be selected before launching the acquisition, and the required spatiotemporal resolution can be selected after.

**Figure 4:**
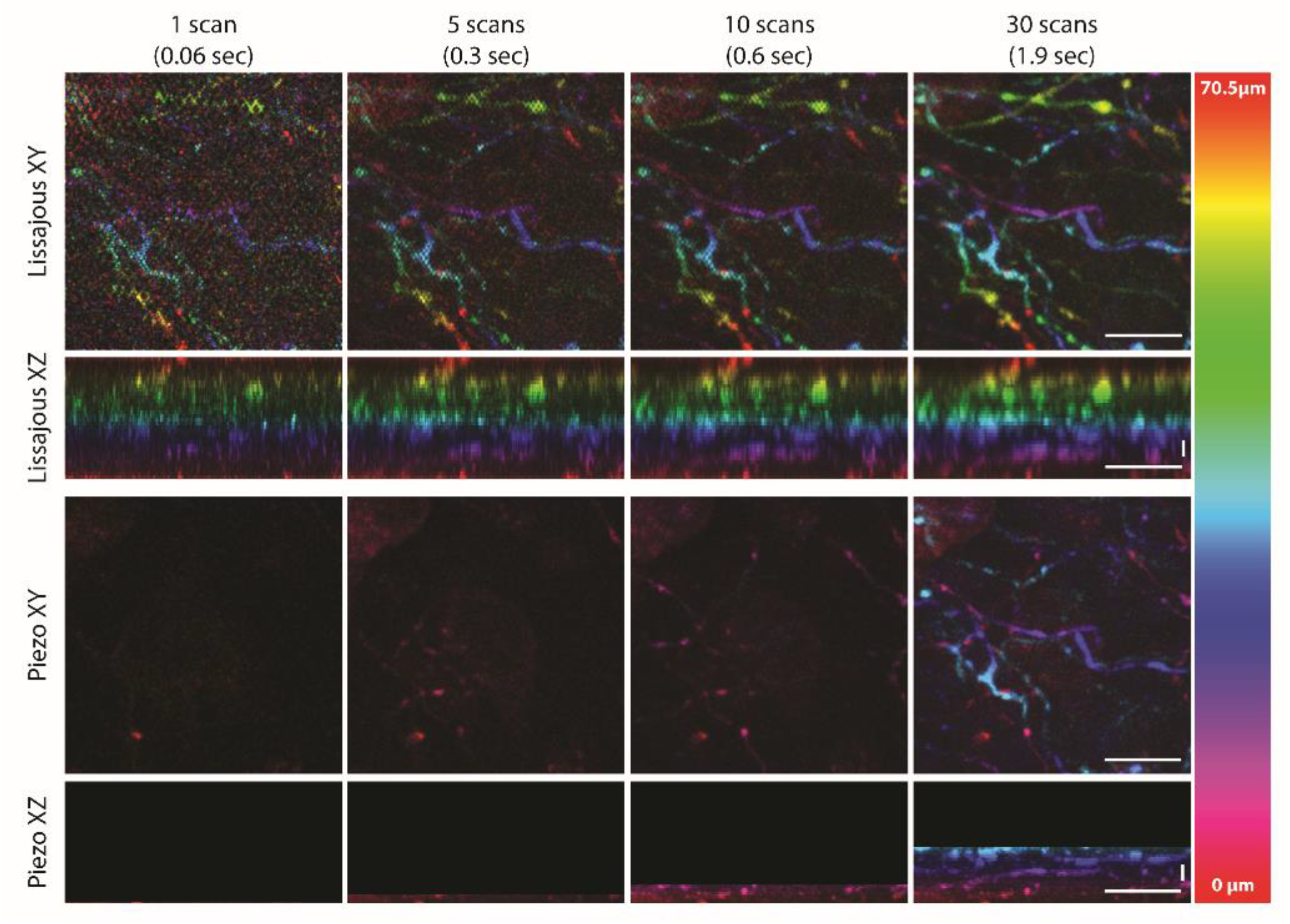
Instant access to volumetric information. Depth-color coded images of neurons in a brain slice acquired by Lissajous scan (and a standard piezo scan. First and second rows present reconstructed 3D volume acquired by Lissajous scan using 1, 5, 10, and 30 scans, showing that 3D information was obtained even with a single scan, and higher spatial resolution and SNR were achieved with more scans. The third and fourth rows present the images acquired by traditional piezo scanning, showing how 3D information is sequentially acquired one plane at a time. Therefore, even with 30 frames only half of the volume was imaged. Note that the step size of the piezo scan was 1.1 µm, at the Nyquist sampling rate of the objective lens, which required 64 frames to complete the entire axial range. Scale bar 10 µm.

### 3.3 Fast volumetric imaging of calcium transients

Traditional confocal microscopes face several issues when monitoring the dynamics of fast events — their 3D imaging speed might not be sufficient to characterize the process under study, it can be difficult to determine the optimal tradeoff between spatial and temporal resolution and, even if such optimal conditions are found, they may change over time. In these instances, volumetric Lissajous confocal microscopy can offer a competitive advantage. To prove this point we recorded the calcium activity of murine neuronal networks from a 3D cultured *in vitro* brain model. We stained the cells with the fluorophore Fluo-4 AM, which is sensitive to local calcium concentrations (see Methods section). As calcium events have a typical duration of milliseconds and propagate through complex networks distributed across the three dimensions of space, they are extremely challenging to image in commercial confocal systems. Figure 5 shows images and data obtained from a 60 x 30 x 50 µm volume of the brain model over a total of 35 seconds. Each 3D scan took 33 ms, thus capturing a total of 1050 scans. As expected, integrating the information retrieved over multiple scans leads to higher spatial sampling and thus higher resolution (Figure 5a, top row), albeit with an inevitable loss of temporal resolution. Interestingly, it is possible to bypass this problem by using advanced image processing algorithms. Specifically, the information from the non-sampled voxels can be considered as an inpainting problem, where images that exhibit lost or deteriorated parts are reconstructed. Several inpainting methods are currently available, based either on machine learning [20] or compressed sensing [21], among others. In these experiments, we used an iterative model-based inpainting algorithm already used in microscopy [13,22]. As presented in Figure 5a, bottom row, the inpainted image after 1 single scan shows the soma of a neuron and several additional structures distributed over the entire scanned volume with high contrast (arrows in the figure image). Note that from the image counterpart prior to inpainting the soma of the neuron is barely distinguishable. After 3 scans, the inpainted image shows most of the structures that were only visible after averaging 100 scans (Figure 5b). After 5 scans, structural details in the image appeared barely distinguishable when compared to images formed with 100 scans, including the vertical dark lines indicated by arrow heads. To obtain similar results without inpainting, it is necessary to integrate more than 10 scans. Therefore, inpainting offers a feasible route to enhance image quality without sacrificing temporal resolution.

**Figure 5.**
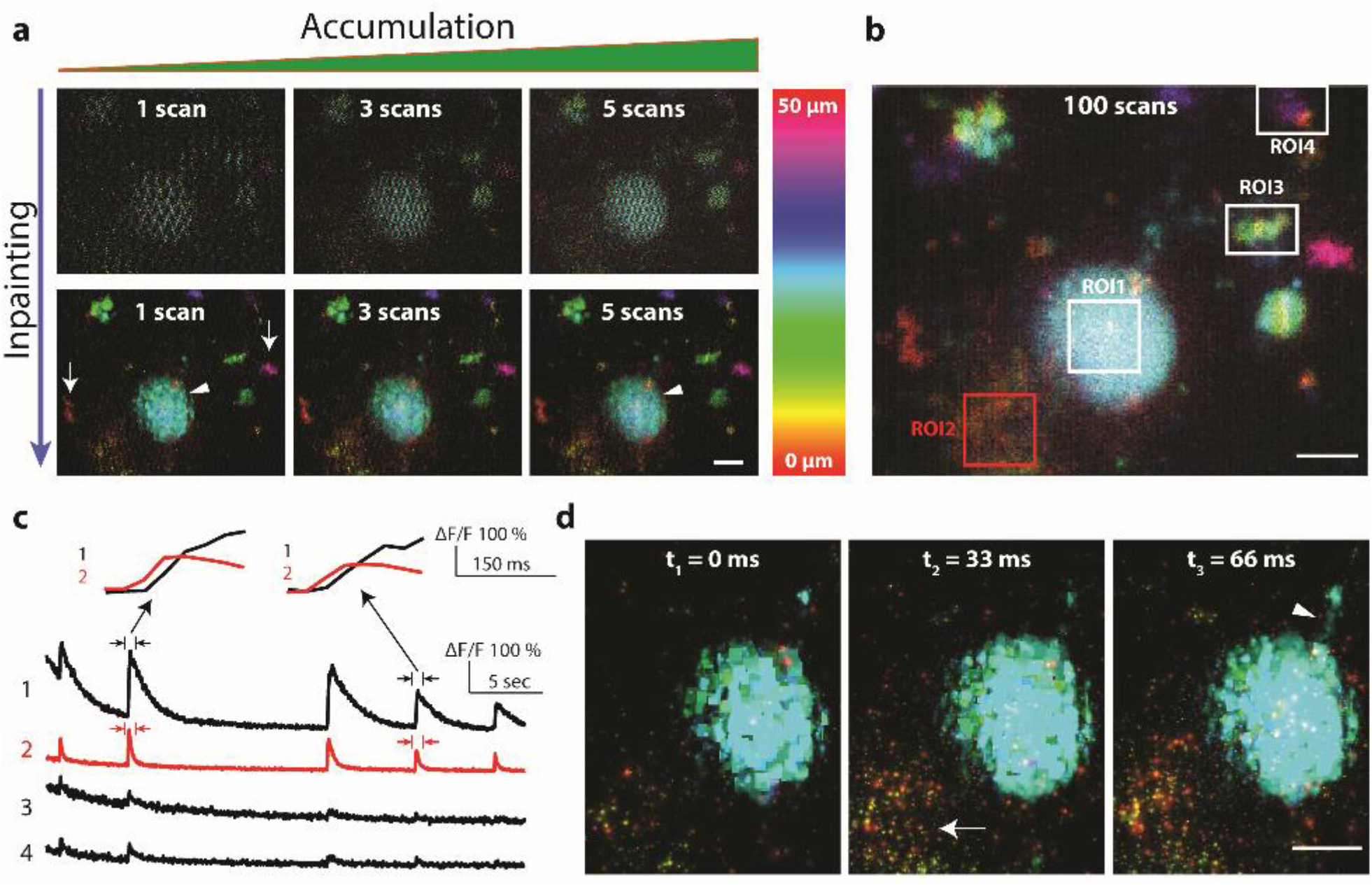
Volumetric imaging of calcium transient in a 3D brain model. a) Depth-color coded images taken by Lissajous scan with time accumulation and inpainting. Integration over multiple scans improves image quality and resolution albeit a loss in temporal resolution (top row). Inpainting algorithms compensate un-scanned voxels and thus the loss of information, improving the image quality (bottom row). b) Depth-color coded image of the brain acquired with 100 scans without inpainting. c) Examples of calcium transients corresponding to the 4 different regions of interest (ROI) highlighted in (b). The intensity plots (ΔF/F) where recorded for a total of 35 seconds, showing typical peaks and subsequent decays caused by neuronal signaling. The magnified plots at two time points for ROI1 & ROI2 show that the signal stared increasing at ROI2 before at ROI1. d) Inpainted images at three consecutive time points with 33 millisecond intervals, taken at T=26 sec. The signal at 0 ms remained relatively low and increased after 33 ms at one particular area (arrow), and then after another 33 milliseconds, the signal increased at the dendrite-like structure (arrowhead). Depth color bar in (b) and (d) is the same as in (a). All images were cropped from the 4D data (a volume of 60 x 30 x 50 µm, for 35 seconds) shown in Movie 2. Scale bar is 5 µm in all images.

In the study of neuronal networks, the dynamics of calcium transients, i.e. local variations in calcium flux which lead to important cellular functions for neuronal development and physiology, are analyzed at different regions over a large volume [23]. Figure 5c shows intensity plots (ΔF/F) at four regions of interests (ROIs) at different z-positions (note the ROIs in Figure 5b)—the peaks and subsequent decays indicate typical variations of local calcium concentrations caused by neuronal signal transmissions. The magnified plots at two time points show that the system time resolution was high enough to discriminate that the fluorescence signal at ROI2 started increasing just before it rose at ROI1. These temporal variations of the calcium concentrations are visualized in Figure 5d where three images acquired at 33 milliseconds intervals show that the signal intensity increases in different regions at different time points. At t1 = 0 ms, the neuron exhibited relatively low signal and at t2 = 33 ms the fluorescence signal increased at the region indicated with an arrow, and at t3 = 66 ms the signal increased at a dendrite-like structure (arrowhead). Such signal variations at different time points were resolved over the entire 3D space (Supplementary Movie 2), and this indicates that Lissajous confocal microscopy may be a powerful tool in the study of functional connectivity of neuronal circuits.

### 3.4 Ultrafast x-z imaging

Our Volumetric Lissajous confocal microscope has the particular feature of having the slow scanning oriented along the y-axis. Even if it were possible to add a resonant mirror for this axis, the expected gain in scanning speed would greatly increase the systems complexity due to the additional synchronization and faster electronics needed. Alternatively, our current configuration reduces 3D scan time by limiting the travel range of the galvo mirror by having less pixels along the y-direction. This original method of scanning could find applications in imaging flow cytometry [24], particle imaging velocimetry [25] or other instances where fast x-z imaging is preferred. As an extreme example, we imaged an object moving along the z-direction with a static galvo mirror, acquiring 2D sections across the x-z plane. Here, the x-z scan was completed in only 62 µs. Figure S5 shows the x-z images obtained of an oscillating mirror characterized at 5300 and 1600 frames per second. Such ultrafast speed enables object displacements at 1.7 mm/s to be distinguished. As such, the initial acceleration of the object, the posterior advance at constant speed, and the final deceleration can be temporally resolved. Arguably, the lower SNR of normal fluorescence samples would impede the use of this speed for bio-imaging. Still, continuous advances toward more sensitive detectors and brighter dyes are increasing the photon budget of optical microscopes, rendering methods for fast beam scanning more relevant than ever [26].

## 4. Conclusions

Combining an ultrasound varifocal lens with a resonant mirror leads to a novel type of confocal microscope where the laser beam is scanned following dynamic Lissajous trajectories across a volume. Continuous interrogation of specimen at sub-microsecond time scales and sub-Nyquist sampling enables minimizing the light dose exposure and optimizing the data content based on application. In this way, instant or accumulated observation of large volumes can be achieved in a post-processing step, resulting in 3D images with a user-selectable spatiotemporal resolution. Additionally, the system preserves all the key features of conventional CLSM and is easy to implement in a commercial microscope.

Optical imaging techniques capable of capturing continuous changes of living specimen at sub-cellular resolution have been a long-term quest in science. Several strategies have been successfully developed, but they typically involve a tradeoff between spatial or temporal resolution. As our results demonstrate, volumetric Lissajous confocal microscopy provides a unique flexibility for selecting, at any given instance, the optimal conditions for characterizing biological phenomena. The new microscope opens the door to monitoring rapidly evolving processes or events that occur over temporally or spatially varying scales. We anticipate that coupling the technique with parallelized detection methods [9] or nonlinear excitation [15, 29] will lead to improved signal-to-noise ratio, spatial resolution or penetration depth, helping to reconstruct complex 3D phenomena with maximal detail.

## 5. Methods

### 5.1 Implementation of the volumetric Lissajous confocal microscope

All experiments used a commercial confocal microscope equipped with a resonant scanning mirror (Nikon A1R, Nikon Instruments, Japan), four excitation continuous wave lasers (405 nm, 488 nm, 561 nm, 640 nm), and four photomultiplier tubes (two normal and two GaAsP PMTs). The typical resonant frequency of the resonant mirror was 7929 Hz. In all experiments, the confocal pinhole was set to 1.5 Airy units (in the case of multi-color excitation, the Airy unit was calculated for the shortest excitation wavelength) and the pixel size was selected to be smaller than the Nyquist sampling rate for each objective lens. Unless otherwise specified, the resonant scan direction was bi-directional. The image size along the x-direction was always set to 1024 pixels (the maximum available for the current system) and the pixel along y-direction was typically 512 pixels but modified for each experiment. To implement the fast axial scanning in this commercial system, we detached the scan head from the microscope body and inserted a pair of relay lenses (f60 mm and f75 mm at f_TAG_ = 142 kHz, and f200 mm and f200mm at f_TAG_ = 142 kHz) between the tube lens and scan lens. Then, we placed a TAG lens (TAG lens 2.0, TAG Optics Inc., USA) between the relay lenses at a conjugate plane of both the scanning mirrors and the back focal plane of the objective lens. The TAG lens served as a resonant axial scanner that, unless indicated, was driven at a frequency of 457 kHz using the driving kit provided by TAG Optics. To avoid aberrations caused by the Bessel-like refractive index profile of the TAG lens [27, 28], the entrance pupil of the TAG lens was physically blocked with a 1.7 mm aperture. Experiments were performed using either a 40x water immersion (Apo LWD 40x WI λS DIC N2, NA 1.15, Nikon Instruments, Japan), a 20x air (Plan Apo VC 20x DIC N2, NA 0.75, Nikon Instruments, Japan), or 60x water (Plan Apo IR 60x WI DIC N2, NA 1.27, Nikon Instruments, Japan) objective lens. The limited size of the TAG lens aperture reduced the beam diameter of the excitation laser, and the filling factor at the back aperture of the objective lens was approx. 60 % for 40x objective and 45% for 20x objective lens. This problem could have been solved by placing the TAG lens before the scan head and expanding the beam diameter [15,29], but in our current commercial microscope this was not possible. The magnification factor caused by the difference of focal length between the relay lenses caused a smaller field of view, with the pixel size calibrated accordingly. When the TAG lens is off, the microscope operates as a standard confocal system. In this case, we used a piezo scanner (P-736 ZRN, Physik Instrumente, Germany) for axial sample translation.

### 5.2 Image acquisition and processing

To reconstruct an image it is necessary to know the x-y-z position of the focused beam at any time instance. This normally requires a fast acquisition card. Instead, we designed a scheme that obviates the need of fast electronics and can be used with the traditional acquisition card of a commercial confocal microscope. We sent the clock signal generated by the TAG lens driver (adjusted to indicate when the induced refractive index was at a minimum) to the input electronic board of one of the PMT channels (Channel 1), located in the microscope controller box of the Nikon microscope. The commercial software of the microscope generated a black image with some bright pixels corresponding to the TAG lens clock signal (named “reference image” hereafter). Note that such an image corresponded to a Lissajous trajectory, in which each pixel has an associated unknown z-position. We then proceeded to reconstruct the Lissajous trajectory, thus decoding the relationship between the pixel and z-position, using a custom-written Matlab code. For each line, a Lissajous pattern was fitted to match its minima positions to each peak pixel (TAG clock signal) in the reference image (Figure 1d). Once the Lissajous trajectory was reconstructed, the same code enabled sorting the information into a z-stack with an arbitrarily number of axial planes. Importantly, the axial scan range produced by the TAG lens depends on the driving voltage amplitude, frequency or microscope objective used. This value was obtained before starting each experiment by recording the reflected light from a mirror as it was axially translated.

Once the z-stack was retrieved, we processed the images using ImageJ. To improve visualization, we adjusted image contrast, and in some instances we used a Bandpass Filter for removing the vertical lines and a Gaussian Blur for removing noise. Quantitative data analysis was performed on raw images without post-processing. For the measurement of FWHM values of the nanospheres Gaussian fitting was used on the intensity line profile of each nanosphere. Image inpainting was performed using an existing open source Matlab library (Plug and Play priors, available at https://engineering.purdue.edu/~bouman/Plug-and-Play/).

### 5.3 Sample preparation

#### 3D beads sample

The sample consisting of 3D fluorescence nanospheres was prepared by diluting yellow-green 100 nm fluorescent beads (FluoSpheres™ Carboxylate-Modified Microspheres, 0.1 µm F8803, ThermoFisher Scientific, USA) into water at a concentration of 3 × 10^9^ particles/mL. The solution was mixed with 1% agarose solution (Invitrogen Ultrapure Low Melting Point Agarose, Invitrogen Corp., USA), and put on a microscope coverglass.

#### 3-color brain (For Figure 3 & 4 & S6)

The optically-cleared brain slice was prepared following the protocols described in earlier studies [9,30]. Briefly, the tissue was fixed with 4% paraformaldehyde, sectioned to 500μm, labeled with Alexa Fluor® 488 wheat germ agglutinin for the blood vessels, Alexa Fluor® 633 anti-tyrosine hydroxylase antibody mainly labeling the dopaminergic neurons, and SYTOX® Orange labels for the nuclei, and mounted in the RapiClear® 1.52 reagent (SunJin lab Co., Taiwan).

#### 3D neuronal culture for calcium imaging

Primary hippocampal neurons from postnatal C57BL/6 J mice (P0), isolated as previously described [31], were obtained from sacrificed animals respecting the 3R principle, in accordance with the guidelines established by the European Community Council (Directive 2010/63/EU). 3D neuronal cultures in alginate were prepared as already reported [32,33]. Briefly, a cell suspension [107 neurons/mL] in 0.15% (w/v) sodium alginate (Pronova UP LVG, Novamatrix, Norway) was loaded into donut-shaped (3mm inner diameter) 0.8% agarose (Sigma-Aldrich) molds which were surrounded by Neurobasal medium (Invitrogen) supplemented with 10 mM CaCl_2_ (Sigma-Aldrich) to induce alginate gelation. After 30 min, the gelation solution was replaced by Neurobasal medium without additional CaCl_2_. Half of the medium was changed once a week, until calcium imaging was performed. For imaging the spatiotemporal dynamics of the calcium activity, after 21 days, *in vitro* (DIVs) 3D neuronal cultures were loaded with the fluorescent calcium indicator Fluo-4 AM (5 µM) (F14201, Thermo Fisher Scientific), in extracellular recording solution (95 mM NaCl, 5mM KCl, 1.8 mM CaCl_2_, 0.8mM MgCl_2_, 1 mM NaH_2_P0_4_-2H_2_0, 23 mM NaHCO_3_, 10 mM HEPES and 10 mM Glucose (Sigma-Aldrich), pH 7.3), for 15 min. Thereafter, the calcium indicator was washed away and neuron cultures were maintained in the standard recording solution during imaging.

## Funding

NVIDIA Corporation is acknowledged for the GPU grant program (Quadro P6000).

## Author contributions

T.D., P.B, A.D and M.D conceived research. T.D. performed experiments. G.P., M.O., prepared samples. T.D. and M.D. wrote the manuscript, with contributions from all authors.

## Acknowledgements

We thank Dr. Contestabile for sharing the staining protocol of the 3D brain model, SunJin lab Co. for the kind donation of the 3-color brain slice, and Dr. Andrea Barberis, Dr. Tiziana Ravasenga and Dr. Alice Polenghi for the murine hippocampal neurons. We also acknowledge Dr. Vicidomini and Dr. Koho for useful discussions, Marco Scotto for his technical assistance, and Nikon Japan for advice on system modifications. We especially thank Dr. Anthony for English proofreading, and Prof. Bouman for insights into the inpainting algorithm.

