## supplementary text and figures for "Volumetric Lissajous Confocal Microscopy"

### Supplementary information:
### Volumetric Lissajous confocal microscopy

**Takahiro Deguchi[1], Paolo Bianchini[1*], Gemma Palazzolo[2], Michele Oneto[1], Alberto Diaspro[1,3], Martí Duocastella[1*]**

[1]*Nanoscopy & NIC@IIT, Istituto Italiano di Tecnologia, via Morego 30, 16163 Genova, Italy*

[2] *Enhanced Regenerative Medicine , Istituto Italiano di Tecnologia, via Morego 30, 16163 Genova, Italy*

[3] *Dipartimento di Fisica, Universita di Genova, Via Dodecaneso 33, 16146, Genoa, Italy*

*

#### Lissajous trajectory length in xz plane

Combining a resonant mirror (x-axis) with the TAG lens (z-axis) enables translating the laser focus across the xz plane. Because the horizontal scanned range (Δx) is fixed, the overall length of the trajectory depends on the frequency and axial scan range of the TAG lens (Δz). As shown in Figure S1 (red line), when the TAG lens is switched-off the scan trajectory corresponds to a single line in the xz plane –the focus remains at the native focal plane of the lens and the trajectory length is simply (Δx). When the TAG lens is switched-on, the trajectory length increases due to the continuous axial focus translation. Specifically, the trajectory forms a Lissajous pattern according to the system of parametric equations described in Figure 1a. This effect significantly increases the number of voxels scanned per one stroke of a resonant mirror, and consequently, scanning speed. At conditions herein, up to a 45-fold increase in the trajectory length was attained.

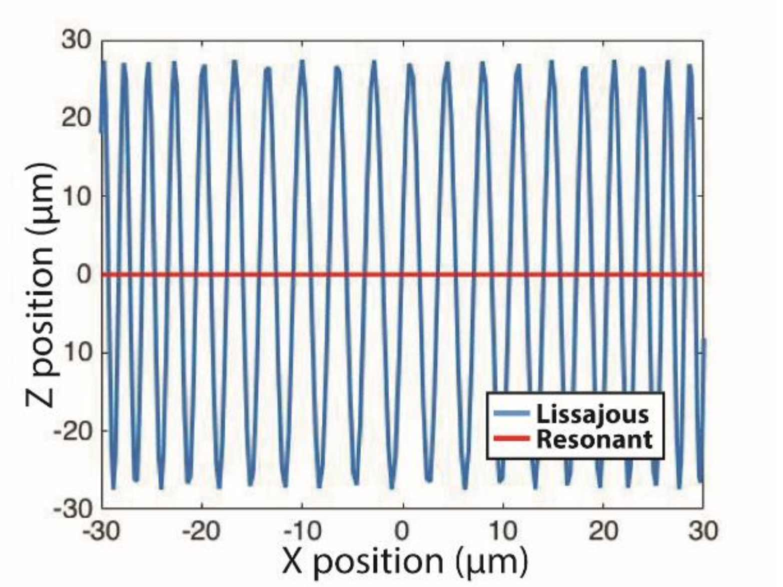


Figure S1: Simulated Lissajous trajectory length in the xz plane. The red line corresponds to the scanned trajectory when the TAG lens is switched-off and the resonant mirror is driven at 7.9 kHz over Δx = 60 µm. Keeping the same x-scanning conditions but with the TAG lens on (frequency of 457 kHz and Δz = 60 µm) results in the Lissajous pattern represented by the blue line. Notably, the trajectory length for the Lissajous scan is 45-fold longer. The graph shows only the field of view of our Nikon A1R confocal microscope, which corresponds to approx. 86 % of the entire stroke of the resonant mirror.

#### Volumetric imaging beyond the typical piezo range

Typical piezo stages used in commercial microscopes cover a scan range of 100 – 200 µm. When the depth of the imaged sample exceeds this range, one is forced to use an alternative translation method, which is typically a linear mechanical actuator that moves the objective lens. This alternative is often slower than the piezo stage, and thus the advantage of Lissajous scan becomes more noticeable. Figure S2 shows images of blood vessels from a brain slice captured with a 20x objective lens and over an axial range of 215 µm. The acquisition by our Lissajous scan (top row) took approx. 2 seconds, while the acquisition with the objective translation (bottom row) took 30 seconds. which is 15-fold longer than the Lissajous scan.

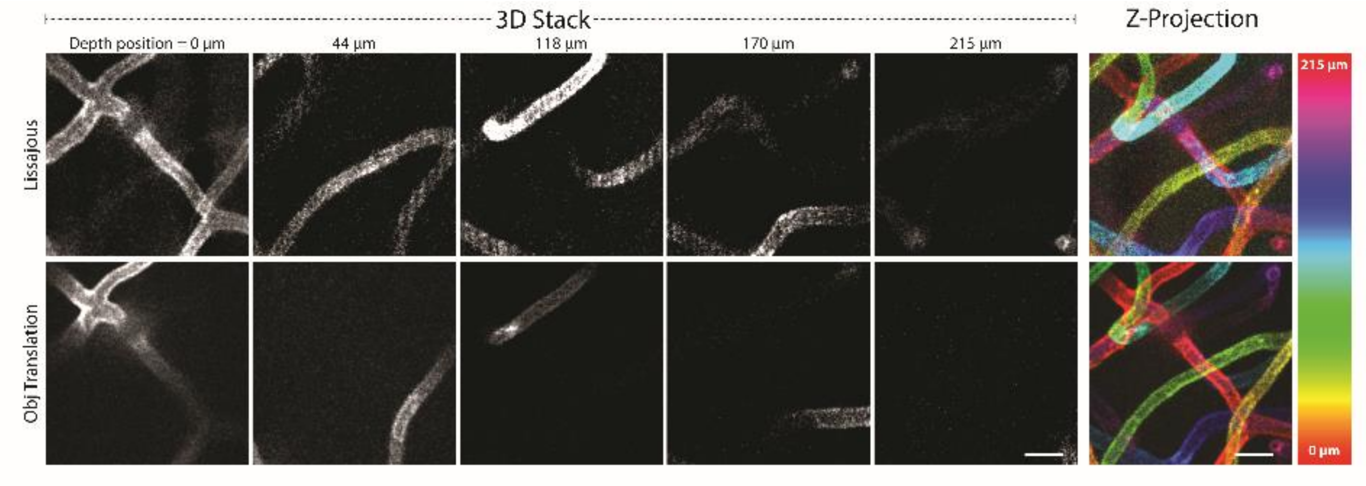


Figure S2: Volumetric imaging beyond the typical piezo range. Images of blood vessels in a mouse brain slice at 5 different axial positions are shown for the Lissajous scan (top row) and the objective translation scan (bottom row). Both stacks were reconstructed with 47 scans, in which the axial step size for the objective translation was set for 4.6 µm. The axial resolution of the 20x objective was 9.2 µm due to the under filling at the back aperture of the objective lens, and thus the step size satisfied the Nyquist sampling rate. The reconstructed volumes are shown in z-projection images with depth-color coding and in supplementary movie (Movie 1). Scale bar 10 µm. 53 x 53 x 215 µm

#### Optical sectioning capability over different axial scan ranges

The optical sectioning of the Lissajous microscope depends on the number of time windows into which the information is divided. To maintain an optimal sampling, the number of time windows should be selected based on the axial scanned range (Δz) or TAG lens frequency ($f_{TAG}$). However, in our current implementation the number of time windows is fixed. Therefore, the effective voxel size in our system depends on Δz and $f_{TAG}$, as shown in Figure S4a. When Δz is large, the available time windows are not enough to sample the entire Δz at the Nyquist frequency. Figure S4b shows a comparison of the axial PSFs in the xz-plane at the same imaging conditions (40x obj, 457 kHz), but with different Δz values. When Δz = 18 µm, the full width at half maximum (FWHM) of the axial PSF was 3.4 µm, close to the value of the piezo scan. The FWHM values increase with Δz — for a scan range of 24 µm, 32 µm, 59 µm, the axial FWHMs were 3.9, 4.5, and 6.7 µm, respectively. Note that the TAG axial scan is continuous and thus each optical slice in a reconstructed 3D stack is a projection of a small volume, so we do not lose information of the sample along axial direction. This contrasts with standard piezo scanning, in which increasing the step size and reducing the axial sampling rate results in some of the planes being skipped and consequent loss of information.

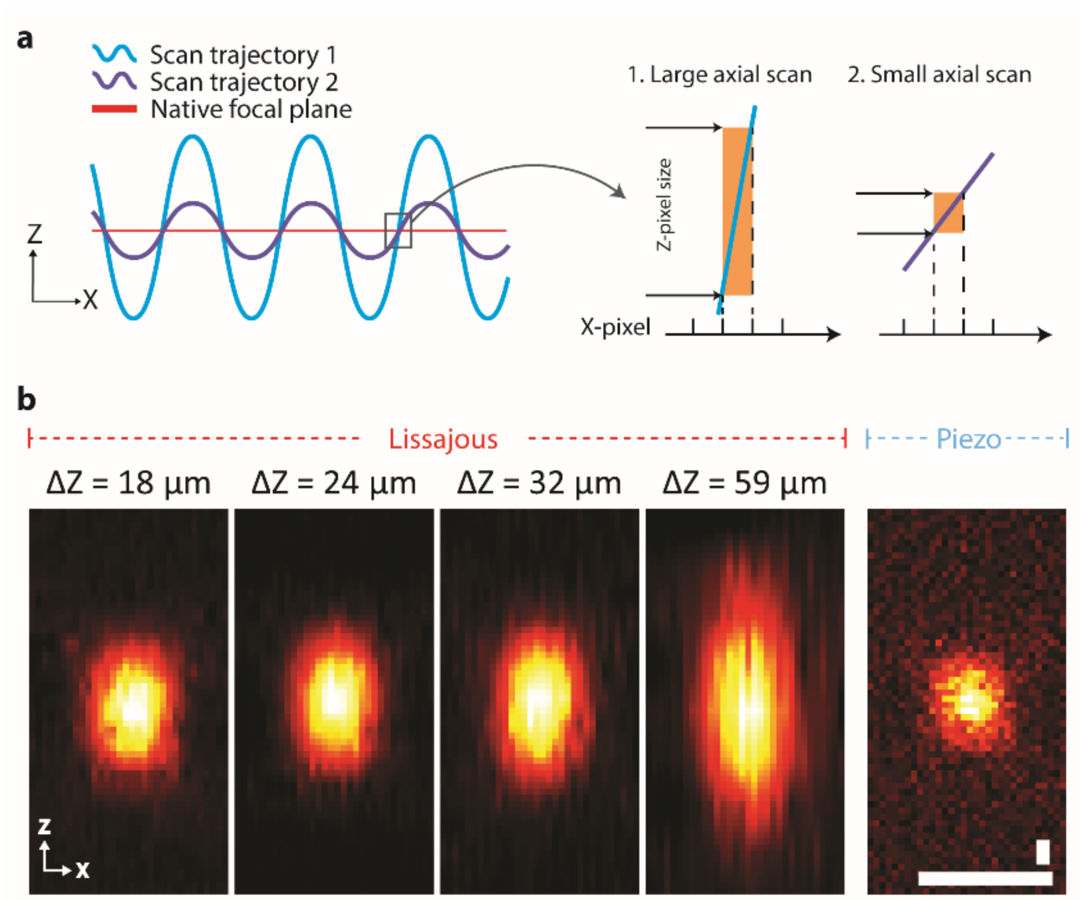


Figure S3: Optical sectioning capability over different axial scan ranges. a) A schematic image of the Lissajous scan trajectories with different axial scan range (Δz) and corresponding pixel size in the z-direction. b) A comparison of axial PSFs between piezo and Lissajous scans with different Δz values. In the case of Lissajous scanning, as Δz decreases, the axial extent of the PSF reaches a value close to that of obtained with the piezo scan. Scale bar 1 µm.

#### Lissajous imaging with increased number of time windows

In order to increase the number of time windows and thus improve the optical sectioning, one can simply reduce the oscillation frequency of the TAG lens or the axial scanned range. This decreases the number of the extrema per a fixed number of time windows, resulting in a higher number of time windows per extrema. Figure S4a shows images of a fluorescent nanosphere (left most) acquired by Lissajous scanning with $\Delta z = 9$ µm, $f_{TAG} = 142$ kHz, and a 60x objective lens (NA1.27), and its image counterpart acquired by the piezo scan. The lateral and axial FWHMs of the PSF were 0.37 µm and 1.5 µm by Lissajous scanning, while the corresponding values by the piezo scan were 0.34 µm and 1.4 µm, respectively. The values of the two axial FWHMs are in a good agreement, confirming that with a sufficient number of time windows the optical resolution of our strategy is not degraded. The images in Figure S4b provide further supports on this—a 3D stack of a flower pollen at five different z-positions show that optical sectioning by Lissajous scanning (top row) is identical to that achieved with a traditional piezo scan (bottom row).

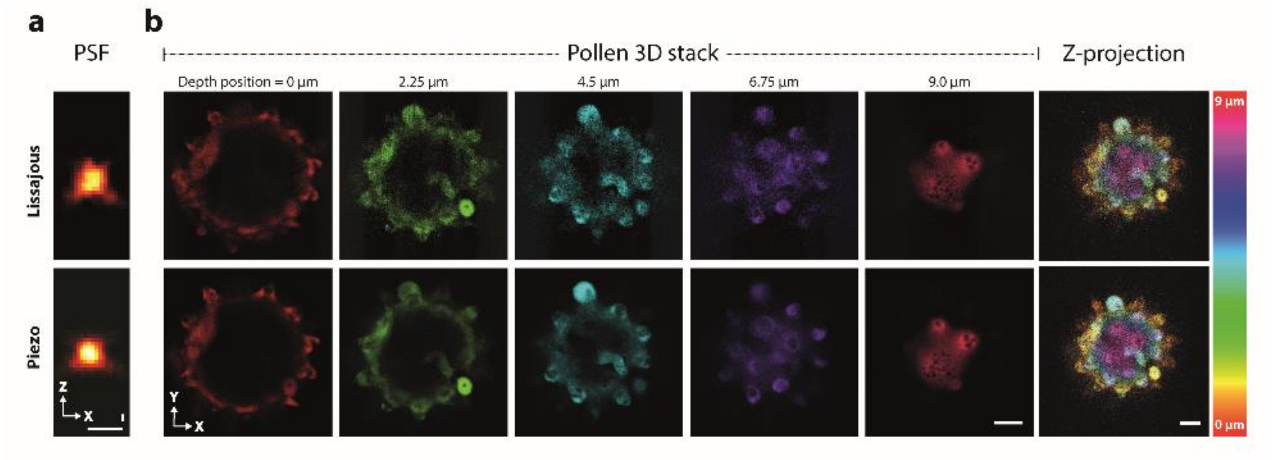


Figure S4: Images with increased number of time windows. a) Top: PSF obtained with the Lissajous scanning while driving the TAG lens at a frequency of only 142 kHz, with an axial scan range of 9 µm, and a 60x objective lens. Bottom: PSF with the TAG lens off (conventional confocal with a piezo stage). b) Images of a flower pollen, acquired with the same imaging conditions, at 6 different axial positions are shown with color coding for depth information. Note that the acquired 3D stacks were reconstructed using 36 scans. Scale bar 5 µm.

### Fast imaging of a moving object

We imaged a fast-moving silicon mirror to demonstrate fast cross-sectional imaging. We placed the mirror on the microscope coverglass and soaked it in water. Using the piezo stage scanner controlled separately from the microscope image acquisition, the mirror was translated along z-axis while we acquired a sequence of images, namely a time-lapse. Figure S5a with a single scan demonstrates how the inpainting process compensates non-sampled voxels.

In Figure 5b, the inpainted time series, exhibiting frame rate of 5.3 kHz and 1.6 kHz frames per second, are shown at three different time points. Such high speed imaging allows distinguishing axial positions of the moving mirror (Movie 4 & 5), resolving its velocity changes over time — the plot of the mirror positions showed three different regions in the velocity, that are an initial acceleration region, a cruising region at constant speed, and a deceleration region (Figure 4c). By fitting the curve with a polynomial function, we obtained the mirror translation velocity of 1.23 mm/sec at the cruising region.

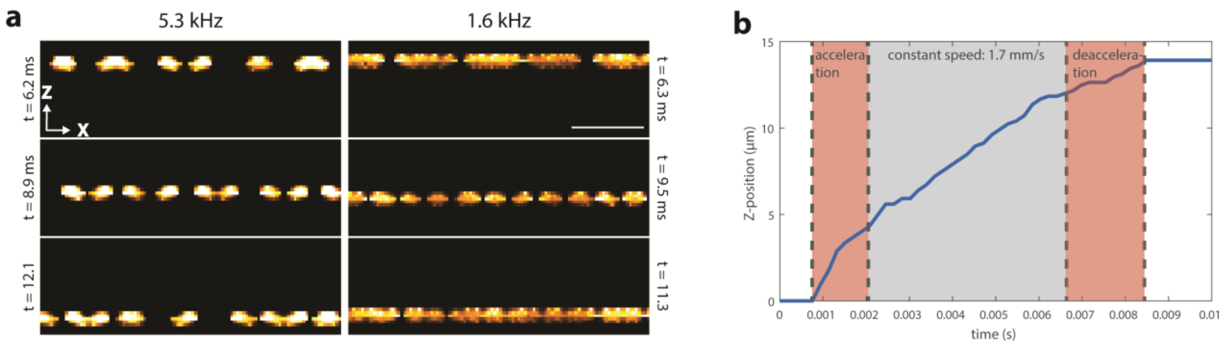


Figure S5. Cross-sectional imaging of a fast moving object. a) A reconstructed cross-sectional image of a moving mirror at 16 kHz (single scan) with and without inpainting (Raw and Inpainted, respectively) and the corresponding magnified images. b) Inpainted images at 5.3 kHz and 1.6 kHz (3 scan and 10 scan accumulations, respectively) at three different time points, showing the axial movement of the oscillating mirror (Movie 4 and Movie 5). The axial scan range was 12 µm and the mirror translation distance was 10 µm. c) A plot of the axial position of the mirror over time (blue) and a fitted curve. Three regions, initial acceleration, cruising at constant speed, and deceleration, with different curve slopes are highlighted. Scale bar 1 µm.
